# The mechanism underlying J-waves and T-waves in the electrocardiogram of mice and zebra finches

**DOI:** 10.1101/2020.07.20.211763

**Authors:** Joost A. Offerhaus, Peter C. Snelderwaard, Jaeike W. Faber, Katharina Riebe, Bjarke Jensen, Bas J. Boukens

## Abstract

Brief cardiac cycles are required to achieve high heart rates as seen in endothermic animals. A main determinant of the cardiac cycle is the repolarization phase of the cardiac action potential, which is visible in the ECG as a T-wave. In mammals with high heart rates – such as rodents – the repolarization phase is short and the ECG is characterized by a positive deflection following the QRS-complex, the J-wave. It is unclear whether birds with high heart rates show similar ECG characteristics. Here we study cardiac repolarization and the ECG in the zebra finch which has high heart rates. In *ex vivo* hearts of zebra finch (N=5) and mouse (N=5), pseudo-ECGs and optical action potentials were measured. In both species, total ventricular activation was fast with QRS durations shorter than 10ms. Ventricular activation progressed from the left to the right ventricle in zebra finch whereas the activation pattern was apex-to-base in mouse. In both species, phase 1 early repolarization followed the activation front, causing a positive J-wave in the pseudo-ECG. In zebra finch, late repolarization was directed from the right ventricle to the left ventricle, whereas late repolarization was directed opposite in mouse. Accordingly, on the zebra finch ECG, the J-wave and the T-wave have the same direction, whereas in the mouse the J-wave and the T-wave are discordant. Our findings demonstrate early repolarization and the associated J-wave are not restricted to mammals and that they also occur within birds. Early repolarization may have evolved by convergence in association with high heart rates.

**Summary statement:** Zebra finches are small birds with high heart rates. Similar to small rodents, the zebra finch ECG contains a J-wave, which is caused by early repolarization

## Introduction

High heart rates distinguish the hearts of mammals and birds from the hearts of ectothermic vertebrates (Hillman and Hedrick, 2015). The high heart rates are required to drive the great cardiac output required to sustain the highly energetically demanding state of endothermy (Crossley *et al*., 2016; Boukens *et al*., 2019). To sustain high heart rates brief cardiac cycles are essential. The main determinant of the cardiac cycle is the repolarization phase of the cardiac action potential. During the repolarization phase, the cardiomyocyte is refractory or unexcitable, which is necessary to reset the intracellular calcium homeostasis. Regional differences in duration of the repolarization phase cause the formation of a T-wave on the electrocardiogram (ECG). The QT-interval is thereby an estimate for the duration of repolarization. Across mammalian species, repolarization differences exist (Durrer *et al*., 1970; Boukens *et al*., 2013; Opthof *et al*., 2017). For example, compared to humans, the action potential of mice is much shorter. In addition, the phase-1 repolarization is large and the plateau phase is absent. Consequently, the mouse ECG is without the isoelectric ST-segment that in humans coincides with the plateau phase of the action potential. Instead, the murine early repolarization is visible on the ECG as a positive deflection directly following the QRS complex (Opthof, 2000). This so-called J-wave is seen in many species of rodents and is therefore sometimes also referred to as the rodent J-wave.

Not all rodent ECGs, however, exhibit a J-wave and the Capybara, the largest rodent on earth, exemplifies this (Szabuniewicz *et al*., 2010). Additionally, in other small non-rodent mammals, such as shrews and bats, J-waves have also been found (Nagel, 1986; Currie, 2018). Despite the difference in phylogenetic origin, these small mammals all share high heart rates that drive the great cardiac output required to sustain their greater mass-specific metabolic rates (Lillywhite, Zippel and Farrell, 1999; Burggren, Farrell and Lillywhite, 2011). In order to maintain a high heart rate, cardiomyocytes must be excitable when the next cardiac cycle arrives. For this, shortening of the action potential is crucial in order to overcome refractoriness and thereby ensuring excitability. Therefore, the cardiomyocyte must adapt by repolarizing fast. This phenomenon is also reflected by the negative relationship between the QT interval on the ECG and heart rate (Bazett, 1920).

If early repolarization is an adaptation to high heart rates, rather than a rodent specific trait, we also expected to find it in the hearts of other endothermic animals with high heart rates (Lillywhite, Zippel and Farrell, 1999). A variety of studies have looked at the electrocardiographic characteristics of different bird species (Table 1). From these studies, not all bird species exhibit a clear isoelectric ST segment, which can be due to early repolarization (Nap, Lumeij and Stokhof, 1992; Cinar *et al*., 1996; Lopez Murcia *et al*., 2005; Szabuniewicz and McCrady, 2010; Hassanpour and Khadem, 2013; Hassanpour *et al*., 2014, 2016). In the absence of recorded action potentials, however, early repolarization has not been shown in any of these studies. In the current study, we electrically characterized the heart of the zebra finch, a small bird with high heart rate (∼600-700 bpm), by recordings of ECG and action potentials and with comparisons to the mouse heart to see if these small birds also show early repolarization (Cooper and Goller, 2006).

**Table 1.** Body mass (kg), heart rate (beats per minute) and ST segment (in lead II) in birds.

| Species | Body mass | Heart rate | Anesthesia | Clear<br>isoelectric<br>ST segment | Reference |
| --- | --- | --- | --- | --- | --- |
| Ostrich | ~100 | 91 | No | Present | (Rezakhani *et al.*, 2007) |
| Emu | 41 | 69 | Yes<sup>#</sup> | Present | (Cushing *et al.*, 2013) |
| Turkey | 14 | 206 | No | Present | (Boulianne *et al.*, 2015) |
| Andean condor | 9.3 | 163 | No | Present | (Wiemeyer *et al.*, 2013) |
| Whooper swan | 9.2 | 85 | No | Present | (Machidad *et al.*, 2001) |
| Griffon vulture | 6.4 | 160 | No | Absent | (Talavera *et al.*, 2008) |
| Pekin duck | 4 | 281 | No | Absent | (Cinar *et al.*, 1996) |
| Green peafowl | 4 | 258 | No | Present | (Hassanpour, Hojjati and Zarei, 2011) |
| Muscovy duck | 3 | 147 | No | Absent | (Hassanpour and Khadem, 2013) |
| Golden eagle | 2 | 347 | No | Absent | (Hassanpour, Moghaddam and Bashi, 2010) |
| Helmeted Guinea Fowl | 1.3 | 338 | No | Absent | (Hassanpour, Zarei and Hojjati, 2011) |
| Peregrine Falcon | 0.80 | 268 | No | Absent | (Rodríguez *et al.*, 2004) |
| Rook | 0.55 | 340 | No | Absent | (Hassanpour *et al.*, 2016) |
| Racing pigeon | 0.52 | 211 | No | Absent | (Lumeij and Stokhof, 1985) |
| Przevalski's partridge | 0.51 | 317 | No | Present | (Liu and Li, 2005) |
| Chukar partridge | 0.42 | 317 | No | Present | (Uzun, Yildiz and Onder, 2004) |
| Pigeon (Spanish Pouter) | 0.35 | 283 | No | Absent | (Lopez Murcia *et al.*, 2005) |
| Teal | 0.30 | 152 | No | Present | (Machidad *et al.*, 2001) |
| Japanese quail (male) | 0.14 | 460 | No | Absent | (Szabuniewicz and McCrady, 2010) |
| Laughing dove | 0.13 | 357 | No | Absent | (Hassanpour *et al.*, 2014) |
| Japanese quail (female) | 0.12 | 320 | No | Absent | (Szabuniewicz and McCrady, 2010) |

<sup>#</sup>Ketamine & Xylazine

## Material and Methods

### Methods

#### Isolation of hearts

Avian subjects were five zebra finches (*Taeniopygia guttata*,) three females (age:), two males (mean age 49.0 days ± 0.8, mean body weight 18.0g, ± 1.1) from the breeding colony at Leiden University. These birds were obtained at three different instances, whenever birds were culled at the colony (independent of this study). After catching the birds were immediately and swiftly killed by cervical dislocation. Immediately following the cervical dislocation, the rib cage was cut laterally and the ventral part was lifted up. The heart was lifted by grapping lung tissue with forceps and the heart was excised by cuts to the lungs, veins, and arteries. At all times, we strove to avoid touching and damaging the heart chambers. The heart was then transferred to a petri dish containing cold cardioplegic solution, containing (mM): NaCl, 110; CaCl_2_, 1.2; KCl, 16; MgCL_2_, 16; NaHCO_3_, 10; and glucose, 9.01 at 4°C. A cannula was inserted in one of the 3 main branches of the aorta and fastened by a suture surrounding all 3 trunks. The arterial pole and coronary vessels were then flushed with ice-cold cardioplegic solution. The procedure from cervical dislocation to the flushing of the coronary vasculature took approximately 5 min. Once the coronary vessels were flushed, excess non-cardiac tissue was trimmed off and the heart was placed in a reservoir of ice-cold cardioplegic solution and kept there during transport from Leiden to Amsterdam (± 45 min).

The use post mortem material of animals culled as breeding surplus is not considered a procedure on itself in accordance with the Experiments on Animals Act (Wod, 2014). This is the applicable legislation in the Netherlands in accordance with the European guidelines (EU directive no. 2010/63/EU) regarding the protection of animals used for scientific purposes. Therefor a license was not obtained for the procedure. All zebra finches were however housed and cared for in accordance to these regulations and internal guidelines concerning care of the animals and licensing and skill of personnel. This also includes that advise is taken from the animal welfare body Leiden to minimize suffering for all animals at the facility (with or without a license).

Five mice (*Mus musculus*) were used (FVB/NRj background, male, mean age 3.3 months, ± 0.1) for the experiments. Mice were kept at the Amsterdam University Medical Center (AUMC) animal breeding unit and had ad libitum access to Teklad 2916 chow (Envigo, Huntingdon, UK) and water. On the morning of the experiment, the mice were moved alive to the department of Experimental Cardiology. Starting the experiment mice were anesthetized by gradually increasing CO_2_ and were sacrificed through cervical dislocation. Mice hearts were excised and the aorta cannulated, in a similar manner as the zebra finch. All mice experimental procedures reported here were in accordance with governmental and institutional guidelines and were approved by the local animal ethics committee of the AUMC.

#### Optical mapping

The isolated zebra finch hearts were transported from Leiden University to the Department of Experimental Cardiology of the Academic Medical Center in Amsterdam. The zebra finch and mouse hearts were mounted on a Langendorff perfusion setup, and perfused at 37°C with Tyrode’s solution (128 mmol/L NaCl, 4.7 mmol/L KCl, 1.45 mmol/L CaCl_2_, 0.6 mmol/L MgCl_2_, 27 mmol/L NaHCO_3_, 0.4 mmol/L NaH_2_PO_4_, and 11 mmol/L glucose (pH maintained at 7.4 by equilibration with a mixture of 95% O2 and 5% CO_2_)), containing blebbistatin to prevent movement artifacts. After a recovery period of ∼10 min the hearts received a bolus injection of 20 μM Di-4 ANEPPS. 3-lead *ex vivo* ECGs were recorded (Biosemi, Amsterdam, the Netherlands; sampling rate 2048 Hz, filtering DC 400 kHz (3 dB)) and analyzed using LabChart Pro (by Mitchells formula). A standard six-lead ECG was calculated as follows: I = L-R, II = F-R, III = F-L, aVR = R-(L+F)/2, aVL = L-(R+F)/2, and aVF = F-(L+R)/2. Activation and repolarization patterns were measured during sinus rhythm and atrial pacing at a basic cycle length of 120 ms (twice the diastolic stimulation threshold).

#### Analysis

Optical signals were analyzed using custom made software (Laughner *et al*., 2012) based on Matlab2018. Local moment of activation was defined as the maximum positive dV/dt of the depolarization phase of the action potential. One zebra finch was excluded from repolarization analysis because of movement artefacts. Repolarization times were taken from 20 and 80 percent of the amplitude of 10 averaged optical signals. Pseudo-ECGs and optical action potentials were simultaneously recorded and aligned by the start of the optical recording. Start of the QRS complex was taken as point zero.

### Statistics

Variables were presented as mean ± s.e.m. and compared using a two-way analysis of variance (repeated factors: phase of repolarization (early or late), and location (left or right ventricle). Activation times, repolarization times, heart rate, PR interval, QRS duration and QT interval of zebra finch and mice were compared using an unpaired t-test.

## Results

### J-waves and T-waves are concordant in zebra finches and discordant in mice

We recorded and compared pseudo-ECGs (pECG) from Langendorff-perfused hearts of zebra finch and mouse (Figure 2). All isolated hearts were spontaneously beating. Table 2 shows the ECG parameters for the zebra finch compared to the Mouse, which were measured as depicted in Figure 1. The average heart rate in zebra finch was higher than that of the mice, but this was not reach statistically significant (476±66 vs. 343±44, *p* = 0.12; Table 2). Figure 2A shows typical examples of the six-lead pECG of the zebra finch (right) compared to the mouse pECG (left). For the zebra finch, the QRS complex was negative in lead I in three animals, positive in one animal and biphasic in one animal. In all mice, the QRS complex was positive in lead I. QRS duration did not differ between species (Table 2). In both species, the QRS complex was directly followed by a positive J-wave advancing into a T-wave that intersected the isoelectric line on average at 65±4.9 ms after the onset of the QRS complex in zebra finches or at 69.9±50 ms in mice. The vector cardiograms in Figure 2C&D show typical examples of the electrical heart axis of the mouse and zebra finch during activation and repolarization. The J-wave (colored blue) was positive in lead I in both species. In contrast, the T-wave (colored red) was concordant in all five zebra finches and discordant in all five mice (although with a low amplitude).

**Table 2.** ECG parameters for the zebra finch and Mouse. \*\**p* < 0.01

| ECG parameter | Zebra<br>finch (n) | Mean<br>± s.e.m. | Mouse<br>(n) | Mean<br>± s.e.m. |
| --- | --- | --- | --- | --- |
| HR (bpm) | 4 | 476±66 | 5 | 343±44 |
| PR (ms)** | 4 | 51.8±5.2 | 5 | 28.1±0.9 |
| QRS (ms) | 5 | 7.7±1.2 | 4 | 8.2±0.4 |
| QT (ms) | 5 | 65.1±4.9 | 5 | 69.9±5.0 |

**Figure 1.**
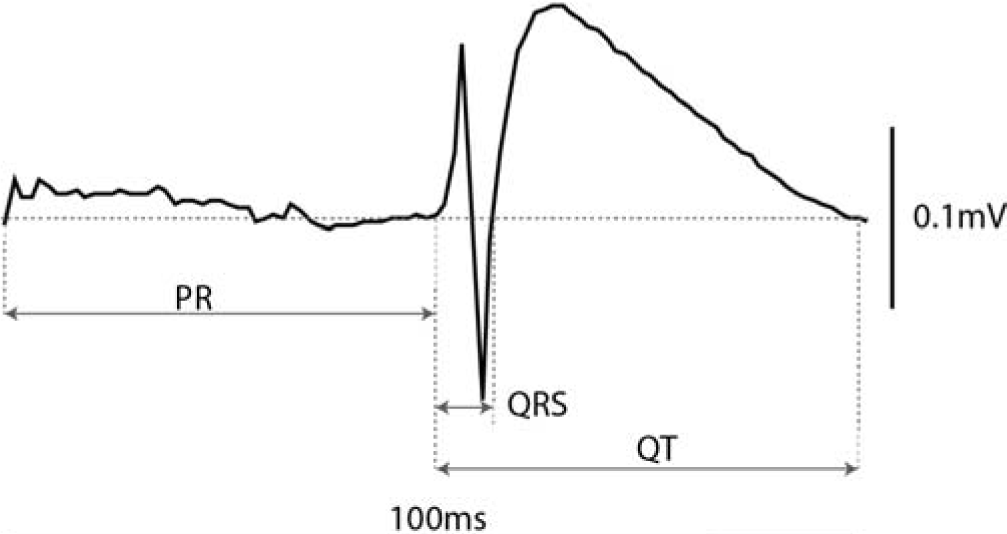
Lead I of the zebra finch ECG showing different ECG parameters.

**Figure 2.**
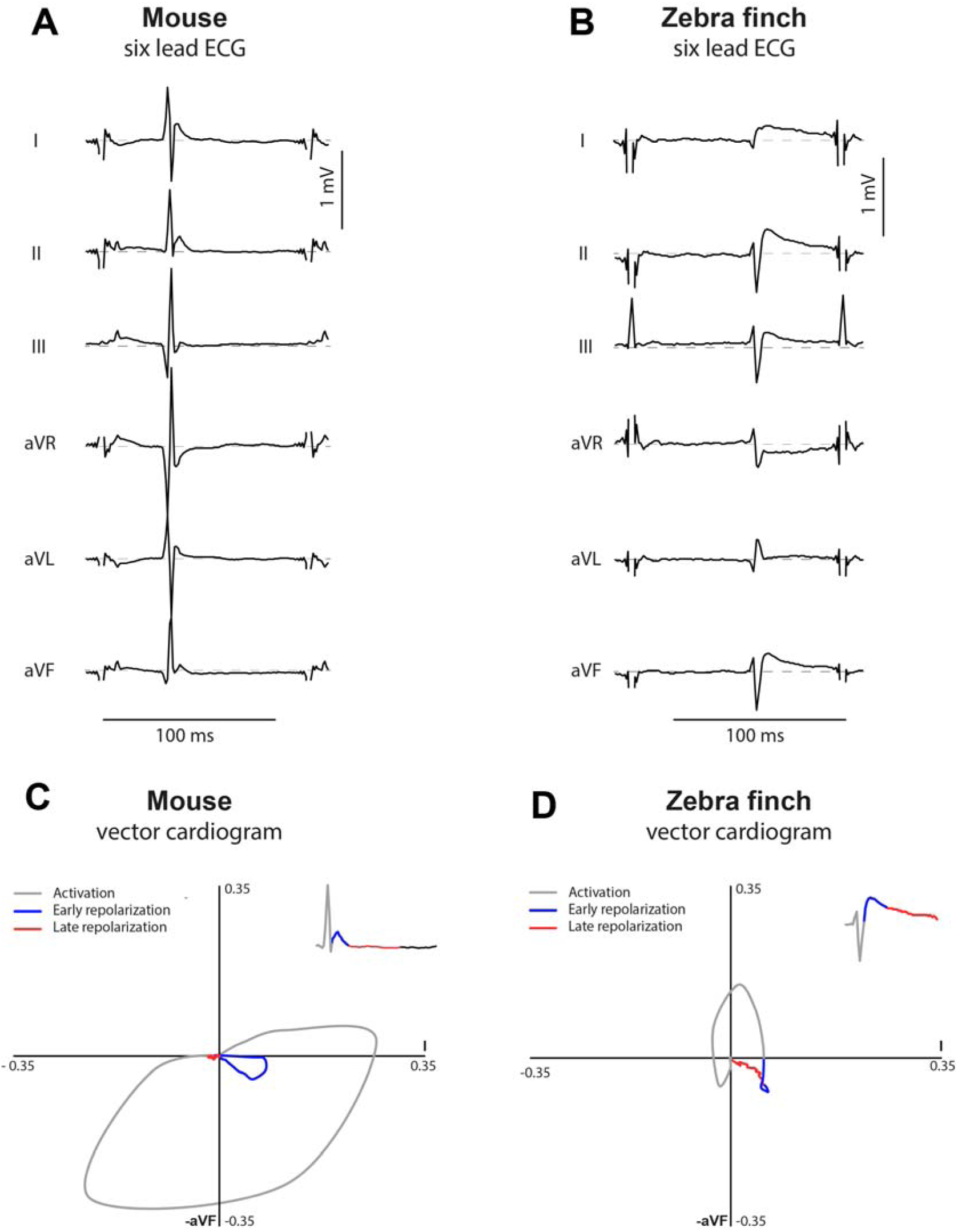
Six-lead ECG and vector cardiograms for mouse and zebra finch: Contrary to mice, the J-wave and T-wave are concordant in the zebra finch. Typical example of a six-lead ECG in the mouse (A) and the zebra finch (B). Both animals show a positive deflection, the J-wave, directly following the QRS. The lower panel shows corresponding vector cardiograms (C and D). The direction of the early and late repolarizing electrical heart axes, characterized by the J-wave and T-wave respectively, are discordant in the mouse (C) and concordant in the zebra finch (D).

### Epicardial activation and repolarization patterns

Concordance between the J-wave and the T-wave is seen when early repolarization as well as late repolarization occur early. To determine the activation and repolarization patterns, we recorded optical action potentials during atrial pacing, in order to exclude an effect of cycle interval on repolarization. Figure 3A shows a typical example of the epicardial activation pattern at the ventral ventricular surface of a zebra finch and a mouse heart. In the zebra finch, the activation front propagated from the left ventricle (LV) to the right ventricle (RV) culminating with the activation of the right ventricular outflow tract (RVOT). In the mouse, the activation front instead propagated from the apex. Epicardial breakthrough of the activation front in the LV occurred earlier in zebra finch than in mouse (0.8±0.3 vs 2.0±0.4 ms, *p* = 0.046 after the start of the QRS complex). However, as with QRS duration, total activation time did not differ between the zebra finch and the mouse (6.0±1.3 vs 5.8± 0.3 ms, *p* = 0.876).

**Figure 3.**
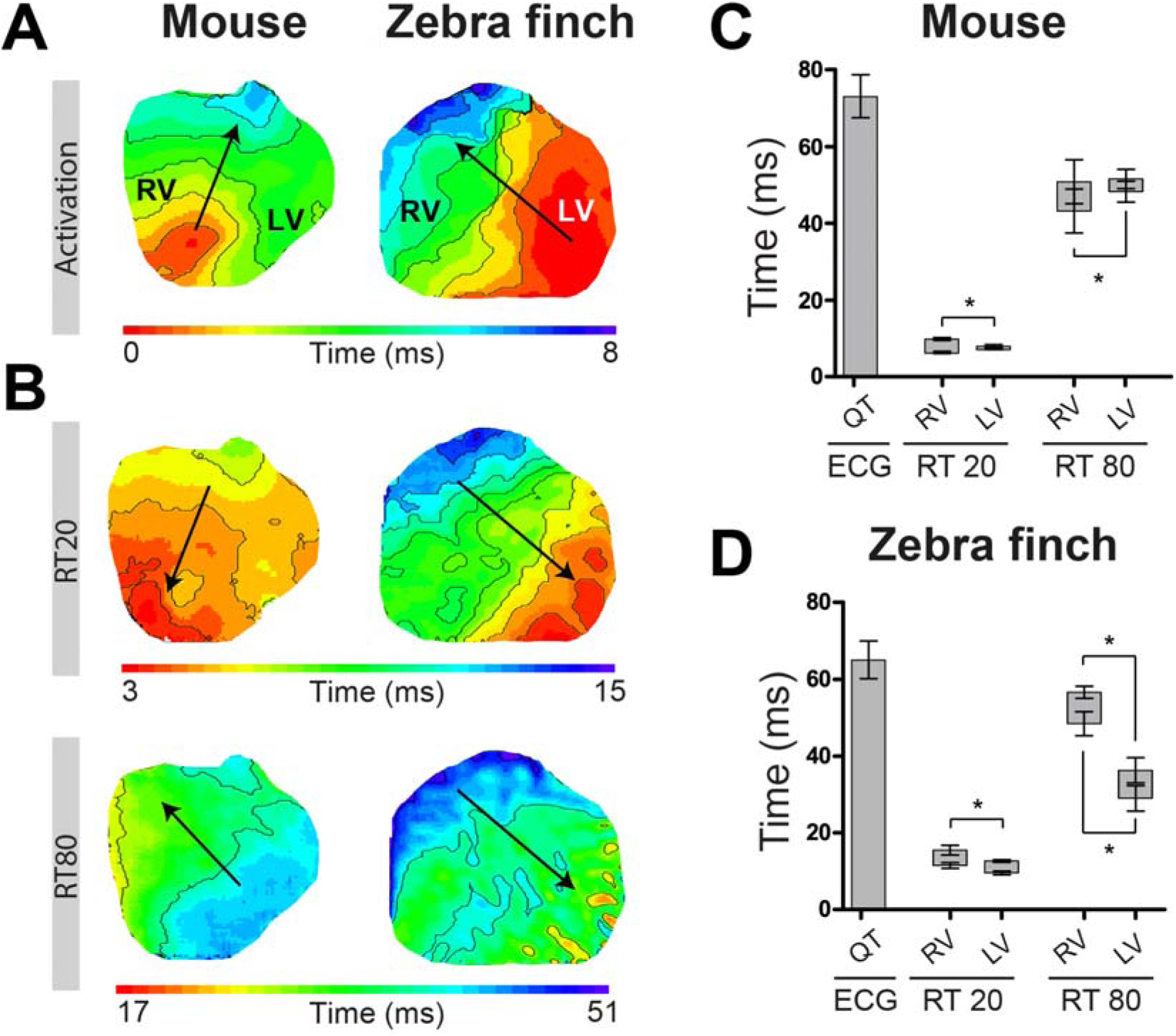
The early- and late repolarization fronts follow the activation front in the zebra finch. Typical activation (A) and repolarization (B) patterns in the mouse (left) and zebra finch (right). The black arrows indicate the electrical vector that is generated by the activation and repolarization front. Early repolarization was defined by 20% of total repolarization (RT20) and late by 80% of total repolarization (RT80). The relationship between early and late repolarization in the right- and left ventricle in mice (C, *n* = 5) and zebra finches (D, *n* = 3) is represented in the bar graphs. RV, right ventricle; LV, left ventricle. \**p* < 0.05.

In both zebra finches and mice, the pattern of early repolarization (20% of repolarization, RT20) followed the activation pattern (Figure 3B). Final early repolarization (RT20) was shorter in mice than in zebra finches (9.9±0.4 ms vs 15.5±1.3 ms, *p* = 0.002). In both species, final early repolarization occurred first in the LV apex and last in the RVOT (15.5±1.3 ms vs 9.5±0.5 ms (p=0.003) in zebra finch; 9.9±0.4 ms vs 7.6±2.2 ms (p=0.041) in mouse). This generated a similar electrical vector, giving rise to the J-wave with similar polarities in the six-lead ECG (Figure 2). During repolarization, an electrical dipole with a negative front in the extracellular space is generated. This is opposite to the positive activation front, as illustrated by the opposed direction of the arrows in Figure 3.

The pattern of late repolarization (RT80) differed markedly between mice and zebra finches (Figure 3B). In zebra finches, late repolarization started 29.0±3.3 ms after onset of the QRS in the LV free wall and ended 56.7±1.6 ms after onset of the QRS in the RV free wall. This pattern generated an electrical vector directed to the left side, leading to a positive T-wave in lead I as observed in the pECG. In mice, however, late repolarization started 43.2±5.7 ms after onset of the QRS in the RV free wall and ended 51.6±2.6 ms after onset of the QRS in the LV free wall. This generated an electrical vector directed towards the right side, leading to a negative T-wave in lead I as observed in the pECG which is opposite to our observations in zebra finches.

Figure 4 shows the relation between local early and late repolarization and the J- and T-wave in zebra finch and mouse. In the zebra finch, both early and late repolarization occur later in the RV compared to the LV, causing a positive J-wave and positive T-wave in lead I. In mice, early repolarization occurs later in the right than LV, whereas late repolarization occurs earlier in the RV compared to the LV (Figure 4A). This latter phenomenon gives rise to a positive J-wave and negative T-wave in lead I.

**Figure 4.**
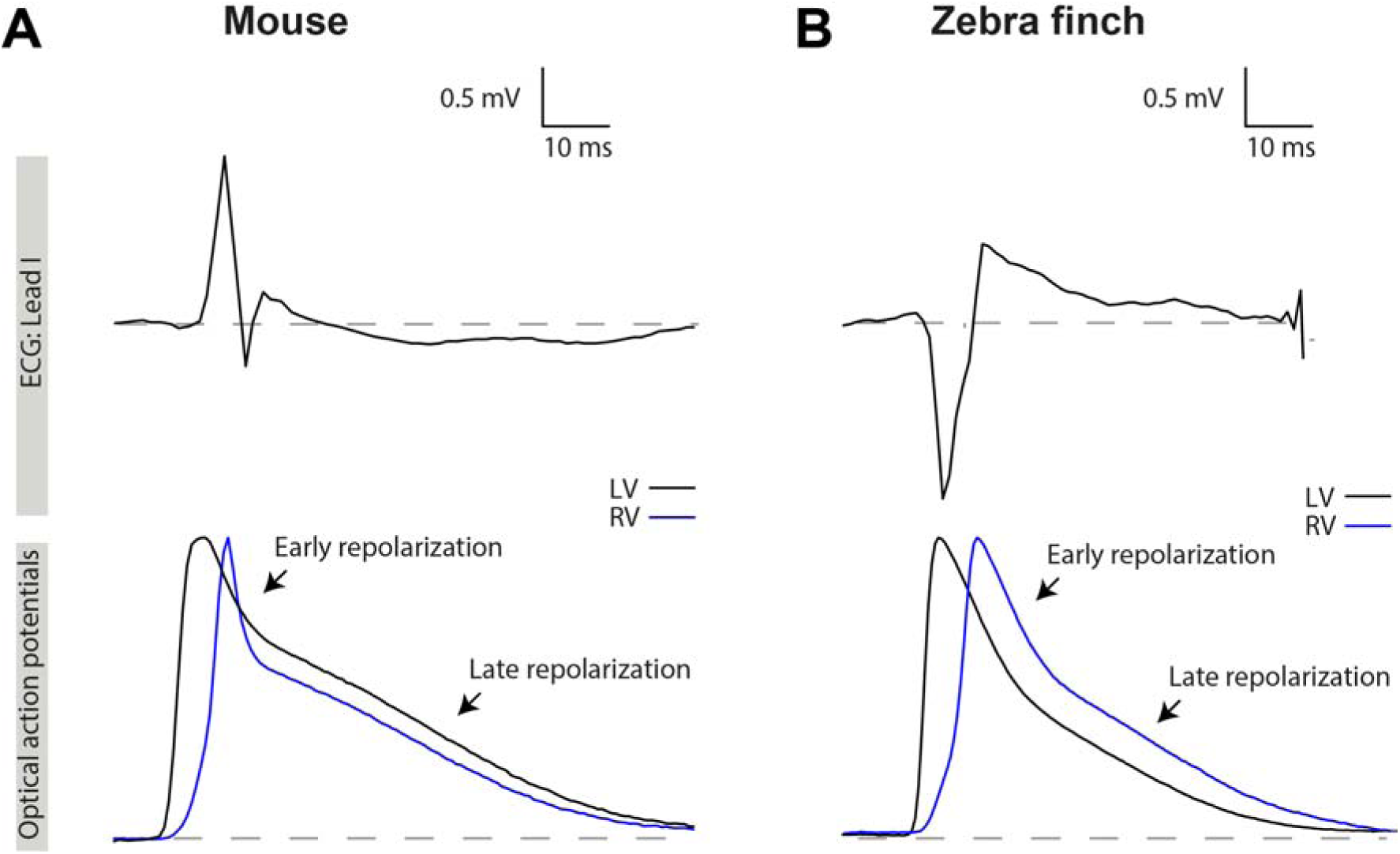
The relation between the J-wave and T-wave results from the order of early and late repolarization. Lead I ECG traces aligned with optical action potentials from the LV (black) and RV (blue) in the mouse (A) and zebra finch (B). The positive J-wave is the result of an early repolarization front moving from LV to RV in both animals. In the zebra finch, late repolarization moves from LV to RV, resulting in a concordant T-wave. In the mouse, however, the wave front moves from RV to LV resulting in a concordant T-wave. LV, left ventricle; RV, right ventricle.

## Discussion

We show that J-waves are not a unique to small mammals, but also occur in at least one bird species. The zebra finch shows a J-wave, which results from early repolarization starting in the LV and ending at the base of the RV. This early repolarization pattern resembles that of mouse, supporting the hypothesis that early repolarization is present in animals with high heart rates. Additionally, we found that late repolarization in the zebra finch heart starts in the LV and ends in the RV leading to a positive T-wave, which is in contrast to mouse, and results in a T-wave with concordance to the J-wave.

Mammals and birds evolved independently from amniotes with hearts that likely were much like the hearts of extant reptiles (Jensen and Christoffels, 2020) and QT duration in mammals and birds is approximately four-fold shorter than in reptiles (Boukens *et al*., 2019). Early repolarization shortens the ventricular action potential and shorter action potentials allow for high heart rates. The presence of early ventricular repolarization in the zebra finch adds to the numerous features of convergent evolution between mammalian and avian hearts (Boukens *et al*., 2019; Kroneman *et al*., 2019). On the level of ion channels, early ventricular repolarization in the mouse is dependent on the currents I_Kur_ and I_to_ (Brouillette *et al*., 2004). It remains to be shown whether early ventricular repolarization in zebra finch is dependent on these same channels. There are multiple species of bird of which the ECG has been published and in which our conjecture that early repolarization of the heart ventricle always occurs in a setting of a high heart rate could be tested. Large birds as the Ostrich and Whooper swan have comparatively low heart rates, an isoelectric ST segment, and no J-wave (Machidad *et al*., 2001; Rezakhani *et al*., 2007). However, the ECGs of many birds studied, especially those with higher heart rates (>100 bpm), do not show a clear isoelectric ST segment and the S deflection is often transitioned directly into the T-wave (Nap, Lumeij and Stokhof, 1992; Machidad *et al*., 2001). We consider it likely that early repolarization contributed to the loss of the isoelectric ST segment, making it likely that early repolarization is a commonly occurring mechanism in birds by which high heart rates can be attained.

The J-wave in the zebra finch and mouse heart mainly results from early repolarization differences (RT20) between the base and the apex. Accordingly, the J-wave was positive in the same leads. The J-wave was discordant with the QRS complex in zebra finches but concordant with the QRS complex in mice (Figure 2). Concordance between QRS complex and J-wave (or T-wave) is caused by myocardium activating early but repolarizing late (Opthof *et al*., 2016). Based on the activation pattern and early repolarization pattern this is the case in both species. Nevertheless, the J-wave was discordant with the QRS complex in the zebra finch. We believe that understanding the polarity of the QRS complex can explain this apparent paradox. Transmural activation has been found to be an important component in determining QRS polarity in mouse (Liu *et al*., 2004). This is illustrated by the positive QRS complex in aVF, indicating activation from base to apex, and the epicardial activation pattern, which is from apex to base. Early repolarization in the septum and subendocardium, which has activated before epicardial breakthrough occurs, starts when the whole ventricle has not yet been activated. Therefore, this early repolarization is obscured by the QRS complex (Boukens *et al*., 2013). The J-wave only becomes visible when activation has ended and the QRS complex disappears. From that moment on, the J-wave only represents the early repolarization difference between the base and the apex. In birds, the discrepancy between the epicardial activating pattern and QRS polarity is less. The first epicardial breakthrough occurs earlier in zebra finches than in mice likely reflecting almost simultaneous transmural activation of the ventricular wall, thereby resulting in discordance between the QRS-complex and the J-wave. Birds are known to have a transmural ventricular conduction system, that could underlie this simultaneous transmural activation (Davies and Francis, 1946; Kharin, Antonova and Shmakov, 2007).

Hearts of mammals and birds have a specialized ventricular conduction system causing fast activation of the ventricles (Davies and Francis, 1946). This is supported by our observations that both total activation time of the heart and QRS duration are not different between the zebra finch and mouse (Table 1). In mammals, distinct early activations on the ventral surface of the left and right ventricle are thought to reveal the presence of the left and right bundle branches of the His-Purkinje system (Sedmera, 2011), but such distinct early activations were not obvious in the zebra finches. Epicardial activation in the zebra finch starts in the left ventricular free wall and propagates towards the RV. The activation pattern is reflected by the negative QRS complex in lead I, which has long been recognized on the bird heart ECG (Whittow, 1999). We have detected similar epicardial activation patterns in hearts of alligators but the QRS duration in these animals is much longer (Jensen *et al*., 2018).

The T-wave was concordant with the J-wave in zebra finch but not in mouse. This is explained by the pattern of late repolarization which is from left to right in the zebra finch and from right to left in the mouse. We think that the direction of late ventricular activation does not play a role in enabling a higher heart rate because both final repolarization and QT time (65.1±4.9 vs 69.9±5.0 ms, *p* = 0.31) are not different between the zebra finch and the mouse. Since activation differences between the ventricles are small, the repolarization pattern is mainly due to differences in action potential duration, which, indeed, was longer in the RV than LV in the zebra finch (49.7±2.3 vs 28.0±3.2 ms, respectively). It is remarkable that in the zebra finch action potential duration is much longer in the RV. In mammals, the action potential is substantially shorter in the myocardium of the RV than in the myocardium of the LV (Volders *et al*., 1999). One explanation for this is the relative lower pulmonary blood pressure, compared to the systemic blood pressure in mammals. However, when birds are compared to mammals, the systemic blood pressure is greater and the pulmonary blood pressure is lower (Whittow, 1999; Seymour and Blaylock, 2000). The left-right difference in ventricular pressure is therefore augmented in birds compared to mammals, yet this is not reflected in the action potential durations.

In humans, the prevalence of J-waves in ECGs varies between 5% and 19% (Offerhaus, Bezzina and Wilde, 2020). In the majority of cases these J-waves are benign. However, in some cases J-waves are a symptom of the early repolarization syndrome and are linked to sudden cardiac arrest due to cardiac arrhythmias (Haissaguerre *et al*., 2008). The Brugada syndrome is another arrhythmogenic pathology characterized by J-waves in the ECG. Together, they are known as J-wave syndromes. It is thought that both syndromes share early repolarization at a cellular level as a common mechanism underlying the J-waves in the ECG and the occurrence of arrhythmias (Antzelevitch *et al*., 2016), although this has been debated (Boukens, *in press*; Hoogendijk *et al*., 2010; Boukens, Opthof and Coronel, 2019). Another occasion when J-waves may occur, is during hypothermia or after cardiac resuscitation which are then referred to as Osborn waves (Osborn, 1953; Jain *et al*., 1990). Whether early repolarization at a cellular level causes these Osborn waves is unclear.

### Limitations

Isolated heart preparations are disconnected from the autonomic nervous system. We cannot exclude that autonomic modulation of repolarization has a role in shaping the J-wave and T-wave *in vivo*. Moreover, it is debated whether the substances required for recording optical action potentials, e.g. di-4-anneps and blebbistatin, affect local activation and repolarization patterns and thereby the shape and duration of the T-waves (Fedorov *et al*., 2007; Larsen *et al*., 2012; Kappadan *et al*., 2020). Nevertheless, a previous study of our group showed that the pECG, recorded in the presence of blebbistatin and di-4-anneps, and the *in vivo* ECG are comparable (Boukens *et al*., 2013). Furthermore, pECGs and optical action potentials were recorded at 37°C, which is 2-4°C below the normal body temperature of the zebra finch, thereby possibly contributing to the J-wave in zebra finches (Skold-Chiriac *et al*., 2015).

## Conclusion

We show that early repolarization in the zebra finch heart causes J-waves in the ECG. This resembles the phenotype in mice and supports the hypothesis that J-waves coincide with higher heart rates. In small birds, early repolarization is likely a commonly occurring mechanism in order to attain high heart rates.

## Author contributions

JAO, BJ and BJB have performed the experiments and written the manuscript. PCS assisted with experimental procedures and edited the manuscript. JWF and KR have critically read and edited the manuscript.

